# Simulated complexes formed from a set of postsynaptic proteins suggest a localised effect of a hypomorphic Shank mutation

**DOI:** 10.1101/2023.10.16.562557

**Authors:** Marcell Miski, Áron Weber, Krisztina Fekete-Molnár, Bence Márk Keömley-Horváth, Attila Csikász-Nagy, Zoltán Gáspári

**Author notes:** Email addresses:* (ACSN) (Attila Csikász-Nagy), (ZG) (Zoltán Gáspári).

## Abstract

The postsynaptic density is an elaborate protein network beneath the postsynaptic membrane built up from the same major proteins but differing between synapses in exact composition and organization. Mutations perturbing generally occurring protein:protein interactions in this network can lead to specific effects, and the translation of biochemical effects to the system level can be especially challenging. In this work we use systems biology-based modeling of protein complex distributions in a set of major postsynaptic proteins including either a wild-type or a hypomorphic PDZ mutant Shank protein. Our results suggest that the effect of such mutations is heavily dependent on the overall availability of the protein components of the whole network. We simulated protein complex formation based on experimental data of various cell types with diverse protein abundances. The simulations revealed that mutations interfering with a single protein interaction can lead to protein complex changes only in specific cell types and in specific complexes. Although our system is currently far from a comprehensive description of the PSD, the results suggest a way to interpret specific mutations within a complex framework.

**Graphical Abstract:** 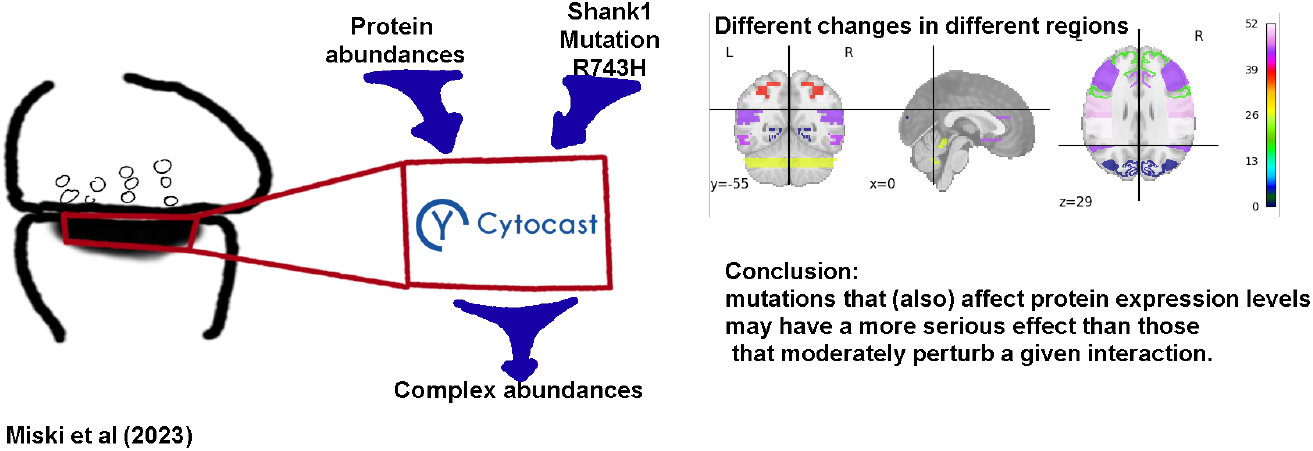

**Highlights:**

- Effect of mutations affecting protein:protein interactions can be modeled in context
- Changes in emerging protein complexes upon mutations are dependent on the availability of the individual proteins
- Our model provides conceptual explanation of confided phenotypic effects of a mutation

## 1. Introduction

The synaptic theory states that the identity of the synapses in different cell types is a key component in establishing the complexity of brain functions, including learning and memory [1, 2]. There are a number of experimental observations indicating that the postsynaptic density (PSD), utilizing the same major constituent proteins, can be highly variable depending on the abundance of its individual protein components [3]. The PSD is also capable of dynamic reorganization during the circadian cycle and upon stimuli [4], and its flexibility to do so has been proposed to play a key role in synaptic plasticity and network rewiring [5]. Despite our knowledge in a number of pairwise protein:protein interactions between postsynaptic proteins [6], the specific large-scale organization of the PSD is still largely elusive.

Mutations identified in proteins of the PSD in various neurological conditions might directly affect specific protein:protein interactions. Although the same proteins and interactions can occur in virtually all postsynaptic compartments, the effect of mutations is often specific to a brain region, to specific interaction partners or leads to the impairment or gain of defined functions instead of leading to the dysfunction of neural transmission in all or most cells [7]. This behavior is expected to be only interpretable by taking into account the complexity of the *in vivo* conditions.

The Shank protein family contains large modular scaffold proteins with both globular and long intrinsically disordered regions [8]. These proteins establish a number of diverse interactions with various postsynaptic proteins. Mutations affecting the availability and/or the structure of Shank proteins have been linked to many conditions [9] from autism spectrum disorder (ASD) [10] to Phelan-McDermid Syndrome (PMS) [11]. In these conditions, Shank3 haploinsufficiency [12], caused by either the complete loss of a copy of the gene or by the presence of a function-affecting mutation, is the most prevalent cause, although mutations in the Shank1 and Shank2 genes can also cause similar, although typically milder, phenotypes.

Besides state-of-the-art microscopic techniques [13, 14], simulation approaches might contribute to our more detailed understanding of the supramolecular structure of the PSD and its changes upon stimuli and mutations. We have previously described extensive simulations on protein complexes using a simplified model of the PSD containing 7 major proteins and have shown that the correspondence between protein component abundance and the distribution of the complexes formed is nontrivial [15]. Specifically, the classification of PSDs based on the abundance of constituent proteins can largely differ from the classification based on complex distributions, a feature supposedly more closely linked to the biological function of the network. Our data set included instances of the same or similar brain regions with different protein abundance, and as such our previous work also models the effect of abundance-affecting mutations. In this work, we address the question of whether hypomorphic mutations, i.e. those reducing but not completely abolishing a function, specifically a binding interaction, can cause measurable effects in our PSD model system. We have chosen the Shank PDZ domain as a model system because of its well-studied nature [16]. Mutations in this domain have been linked to ASD such as R736Q [10] and also have been identified in various cancers [17]. As a model hypomorphic mutation, we have chosen one that can be estimated to cause the decrease of a specific binding interaction between 2and 10-fold. We use the same model system as in our previous work, composed of seven major PSD proteins, except for replacing Shank3 with Shank1 (NMDAR, AMPAR, PSD-95, SynGAP, GKAP, Homer1, Shank1)[15], and more than 500 brain regions with different protein levels [18].

Our results suggest that weakening a single well-defined interaction does not affect the overall distribution of complexes in most investigated brain regions. However, in a small number of cases, the most informative protein complexes defined based on their contribution to the overall diversity of the PSD complexes show a significant change in their abundance. These results indicate that even when the same set of proteins is involved, the biological effect of a mutation can be highly specific depending on the cellular context.

## 2. Materials and Methods

### 2.1 Overview of the simulations, simulation environment

For the simulation of protein complex formation, we have used the agent-based simulation tool Cytocast (cytocast.com). Cytocast provides a modeling platform for examining the diversity of the synaptic protein complexes in terms of the abundance value of their components. Cytocast uses the Gillespie algorithm. Therefore, the number of steps does not linearly scale with the run time of the simulation. The algorithm has to calculate the running time of the next reaction.

The basic principle used by Cytocast also served as the basis for simulations related to Covid-19 [19]. Cytocast and its precursor have been shown to be able to effectively model protein complex distributions even in whole cells [20]. Similarly to our previous study [15], we have investigated a simplified PSD model using only 7 major PSD proteins: AMPAR, NMDAR, PSD-95, Homer1, GKAP, SynGAP, and Shank1. We have replaced Shank3 with Shank1 relative to our previous study because the mutation that was introduced has been described in Shank1 (see below).

Overall, we have simulated protein complexes in 524 different data sets originating from 27 brain region types, as in our previous work[15]. The source data sets contain mRNA expression levels[18] that we have translated into protein abundance[15]. It should be noted that no information about the individuals from which the data are derived is available, whether they can be considered as in a healthy or diseased condition. To estimate the reproducibility of the simulations, 40 simulations were performed for each data set. The raw input abundances can be found in the Supplementary material Appendix A.5. The average copy numbers of the individual proteins, serving as input for the simulations, show a high variance (Table 1).

**Table 1.**
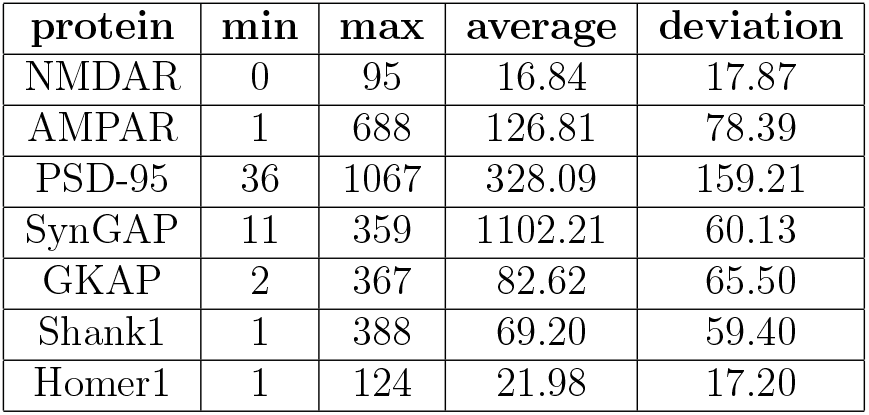
Statistical data on input abundance of proteins derived from mRNA expression levels from article [18]

Dissociation constants were taken from the data described in the literature [21]. In our Cytocast simulations, the dissociation constant was implemented by adjusting the unbinding rate as a result of a mutation, while not changing the binding rates.

### 2.2 Choice and modeling of the Shank1 R743H mutation

In order to use a well-studied interaction, we have chosen the Shank1 PDZ domain and its interaction with the C-terminus of GKAP. The Shank1 PDZ is a globular domain that has been characterized extensively [22]. Moreover, a mutation in the Shank1 PDZ domain has been observed in ASD [10]. The effect of a mutation affecting this region can be easily modeled in our model system of seven major PSD proteins.

In order to model a moderate yet measurable effect of a hypomorphic mutation that weakens but does not abolish binding, we have investigated a number of described mutations for the Shank1 PDZ domain, most of them taken from the COSMIC database [17].

It is not trivial to estimate the perturbation effect of a mutation, especially for one outside the primary binding site, Therefore, we have used the change in domain stability as predicted by the NeEMO method ([23]) and have used this change to estimate the weakening of the interaction by assuming less stable apo and holo structures.

For our modeling purposes, we have chosen the R743H mutation for which we have estimated a 5.5-fold decrease in the binding affinity, modeled as a 5.5fold increase for the dissociation rate of the Shank1 PDZ:GKAP interaction in our setup. This value is regarded as a good compromise between a minimally observable 2-fold change and changing the value by one order of magnitude, a more pronounced change. The arginine affected by this mutation is located on the C-terminal end of helix 2, whereas the R736Q mutation, described in ASD [10], alters an arginine at the N-terminus of the same helix. Both arginines point away from the immediate ligand binding site, thus, both mutations are expected to perturb the binding interaction indirectly.

### 2.3 Analysis of the simulation results: comparing brain regions and identifying the complex with the highest information content in this regard

The identity of protein complexes is not only determined by their composition but also the exact topology, i.e. how the constituent proteins interact with each other, rather also by the binding sites actually participate in the interactions. Complexes with the same protein composition but different binding patterns, which are distinguished during the analysis of the simulation results.

The protein complexes observed in the simulations have been identified and enumerated, and each was assigned a unique identifier. The simulated results for each input data set can be represented by a point in a multidimensional space where each coordinate represents the abundance of a given protein complex.

A protein complex distribution can be described as the linear combination of the emerging protein complexes:

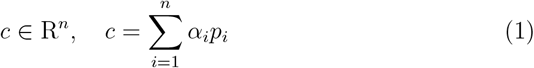

where *α*_*i*_ is the abundance of complex *i* (*p*_*i*_).

Each complex has a characteristic abundance profile across different regions: the complexes with the most varied abundance show the most visible differences between regions. Thus the most variable complexes define the principal components of the system. The principal components are vectors in the previously described vector space, transforming each brain region into the basis of these principal components that allow the largest differences between the brain regions to be visualized.

For each eigenvectors provided by the principal component analysis, the contribution from each complex abundance can be determined. Combining this information with the fraction of overall data variance explained by the eigenvector, it is possible to calculate how relevant and informative the original base vectors (representing complex abundances) are in the comparison of brain regions.

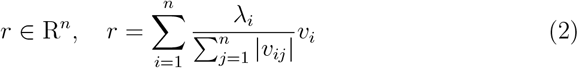

Where *r* is an *n* dimensional vector containing the relevance of each complex, *λ*_*i*_ is the relevance of the *i*-th eigenvector, *v*_*ij*_ is the *j*-th coordinate of the *i*-th eigenvector *v*_*i*_.

### 2.4 Analysis of the simulation results: comparing protein complex abundance

A pairwise T-test is a classical statistic usually chosen when only one change here the mutation is created in the system and the question is how the change affected the mean value. The null hypothesis is that the mean abundance of a complex (in the wild-type and mutant scenario) is the same within a certain level of significance. Thus the alternative hypothesis is that the mean abundance of a given complex formed from the protein set with the wild-type and the mutant proteins differ significantly.

The calculations were performed as implemented in the scipy package (method scipy.stats.ttest rel), where the t-score is calculated by [24]:

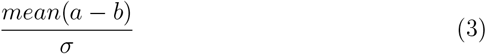

where *a* and *b* are 40-length arrays of the abundances of the given complex for wild-type and mutant simulations respectively and *σ* is the standard error. The p-value is calculated from the t-score based on the alternative hypothesis type that is two-sided:

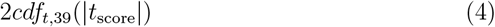

where cdf is the cumulative density function for the T-test with the degree of freedom 39 (for 40 repetitive simulations).

Given that the abundance of each complex is averaged separately during the simulation, the T-test shows which hypothesis we can accept for the abundance only of the given complex. In order to be able to assess the change for all complexes in a specific region, we performed the T-test on each complex and used a weighted average to obtain the P-value. The weights used correspond to the importance of each complex as calculated from the principal component analysis.

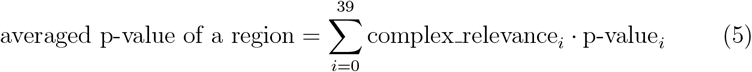

## 3. Results

### 3.1 Identification of the most informative complex

The principal component analyses (PCA) show which complexes are the most informative from the point of view of distinguishing brain regions. The first two principal components for both the wild-type and mutant scenarios cover 44% and 24% of the full variance of the outputs, respectively. The first principal axis is dominated by the abundance of the AMPAR/PSD-95 (id:12) complex, whereas the second one by the PSD-95/SynGAP (id:8) complex.

The most informative complex overall – with the highest contribution considering all principal components and the variance explained by them – is AMPAR/PSD-95/SYNGAP (id:5) with a contribution of 19%. In comparison, the importance of the average complex is very small, approximately

4.48e-06 and the median is 9.98e-08. This is because of the high number of possible complexes.

### 3.2 Overall complex distribution is primarily determined by protein availability

Principal component analysis of the simulation results for the wild-type and mutant scenarios show a very similar overall picture. The two PCA plots can directly be compared as the axes are the same even in the two independent PCA outputs. The data points corresponding to the wild-type and mutant cases move only minimally relative to each other. The average distance between wild-type and mutant regions is 1.5 *±* 0.8.

We note that the PCA does not generally separate the source brain regions (Figure 1). However, the cerebellar-cortex-type regions are well separated from the others.

**Figure 1.**
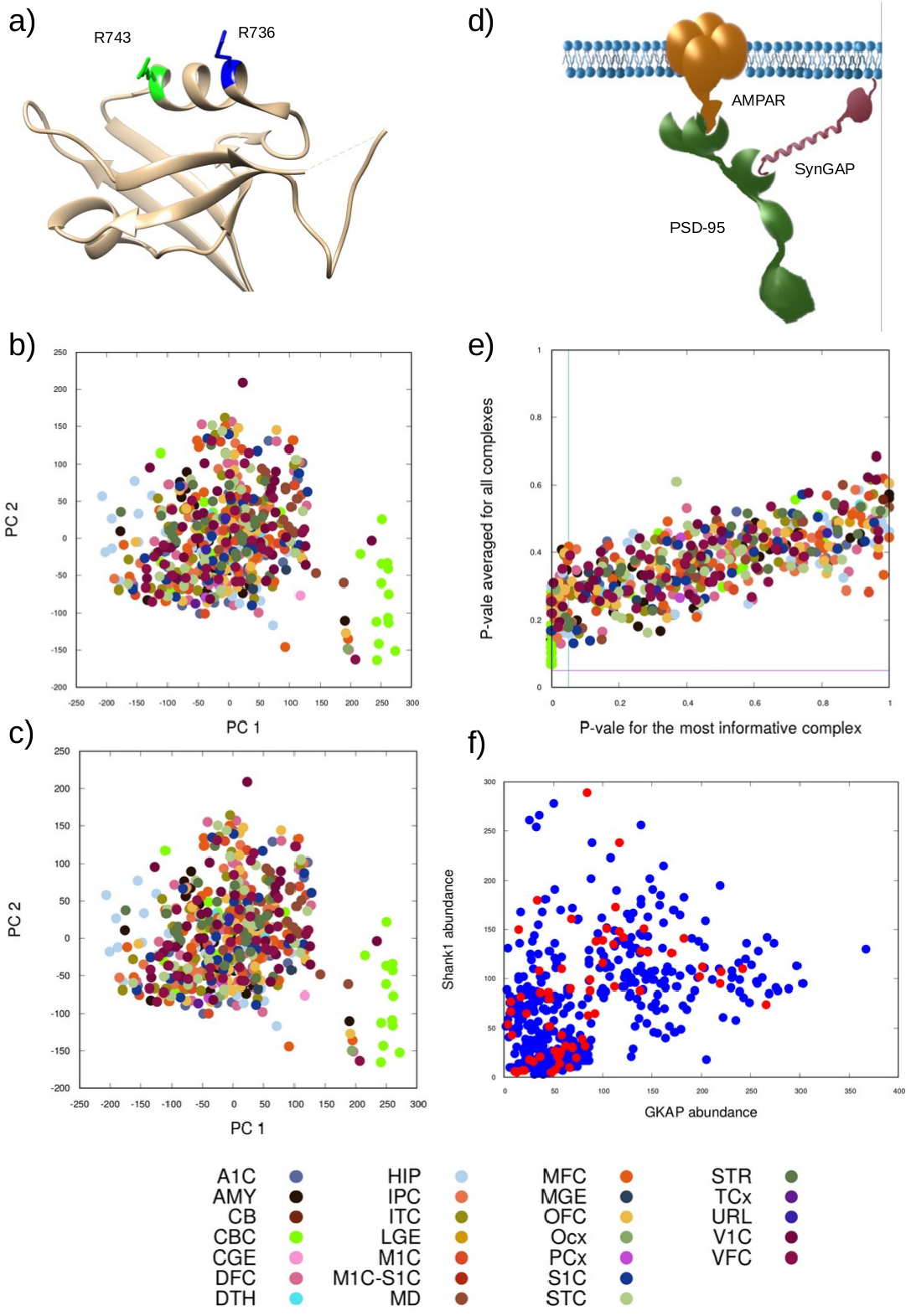
**a)** Position of the mutation selected (R743H, green) and a similar one reported in ASD (R736Q, blue) on the ribbon representation of the Shank1 PZ domain (PDB ID 6YWZ). Both arginines are located on the *α*2 helix flanking the ligand binding groove. Principal component analysis of the obtained protein complex distributions for **b)** the wild-tpye and **c)** the mutant scenarios investigated. Different colors denote different brain regions according to the key at the bottom. **d)** Schematic depiction of the most informative complex according to the PCA (AMPAR/PSD-95/SynGAP). **e)** P-values describing the change upon the mutation relative to the wild-type, the value averaged for all complexes shown vs. those obtained for the most informative complex. The lines denote the 0.05 significance limit. Colors by brain regions according to the key at the bottom. **f)** Abundance of Shank1 and GKAP, the two proteins in the interaction affected by the mutation, in the input data sets. Red circles indicate data sets where the abundance of the most informative complex changed significantly in the output.

Our results suggest that the overall protein complex distribution is determined by the availability of the individual proteins and the presence of a weakening mutation does not cause substantial global effects. This is in line with the system retaining its general functionality. In order to analyze the effect of the mutation in more detail, we have investigated the abundance of the individual protein complexes.

### 3.3 Weakening the Shank:GKAP interaction causes subtle effects in welldefined regions

To analyze the effect of the mutation introduced, we have compared the complex distributions in each region with the wild-type simulations. First, we have compared the cumulative complex distributions using a weighted average of the T-tests conducted for the individual complexes, where the weights were based on the information content (see Methods) of the complexes. These tests show that the cumulative complex distributions remain the same within statistical significance see Figure 1 despite the fact that the dissociative rate of the Shank1-GKAP bond is five times higher in the mutant. Although there is a large variance in the cumulative p-values, each of them remains above 0.05 thus in general for most of the regions and their complexes the null hypothesis is accepted with a significance of 5%. However, the variance in the p-values hints that the regions are not uniformly affected by the mutation, even if the overall change does not reach statistical significance.

However, at the level of individual complexes, there are some exhibiting significant changes in the mutated scenario relative to the wild-type one.

For larger supercomplexes, abundance values above one can be considered more significant than for the smaller complexes due to the much lower probabilities of their formation. Therefore, in the case of supercomplexes, even the smallest appearance is accepted and considered adequate.

Importantly, the p-values of the most informative complex AMPAR/PSD95/SynGAP drop below 0.05 for some of the resulting data sets (Figure 1). This is somewhat surprising as this complex does not contain any of the partners of the interaction affected by the mutation. Thus, it is worth investigating how the abundance of the individual partner proteins Shank1 and GKAP influence the changes observed in the formation of AMPAR/PSD95/SynGAP complex. We have plotted the input abundance of Shank1 and GKAP vs. the p-value of the most informative complex see Figure 1. It is apparent that significant changes are not confined to either the highor the low-abundance regions of the two affected proteins, suggesting that the complex interplay between the interactions of multiple proteins is behind the observed phenomenon.

As a result of mutation, we would expect the complexes in the layer above GKAP, i.e. those containing the membrane receptors and PSD-95, to become more favored since the interaction between GKAP and Shank1 connects these complexes to the larger supercomplexes where Shank1 polymerizes. This effect is observed only in the cerebellar cortex regions and to a small extent. The abundances are shown in Figure Appendix B.1. The complex abundances change from zero to 1 for the complexes SynGAP/PSD95/GKAP (id:9) and AMPAR/PSD-95/GKAP (id:15). However, compared to the other abundances the significance of the appearance of these complexes in the mutant is still very low indicating that the complex abundances mainly remain similar to the wild-type even in the regions with the smallest p-values thus the chosen significance level is acceptable.

Curiously, the regions with the lowest p-values all belong to the cerebellar cortex. The structure and levels of the cerebellum are significantly different compared to the cerebrum, which also means differences in the main neuron types [25]. The differences have already been demonstrated for different Shank3, Shank2 abundances in different layers in the cerebellum [26], and the different aspects of the cerebellum in ASD [27]. It cannot be directly stated that the information appearing in the p-values points to this difference, but it cannot escape our attention.

To get further insight into the changes of complex abundance, we have selected two regions: H376.IIIB.53 M1C-S1C exhibits the lowest nonzero pvalue for the most informative complex (AMPAR/PSD-95/SynGAP) and H376.IV.54 STR has the highest p-value under the significance threshold 0.05 for the same complex (Figure Appendix B.2). In these two regions, the average abundance of the complex AMPAR/PSD-95/SynGAP (id:5) changes from 306 to 301 and from 327 to 321 upon the mutation, respectively. In addition, the abundance of the related complex AMPAR/PSD-95/SynGAP/GKAP increases. Thus, the decrease can be partially attributed to the fact that the complex AMPAR/PSD-95/SynGAP (id:5) associates with GKAP with a higher probability than in the case of the wild-type. The greater availability of uncomplexed GKAP can be explained by the weakened Shank1:GKAP connection.

On the other hand, for the two regions with the highest p-values (H376.XI. 50 HIP and H376.VIII.53 MD), the abundance of the complex AMPAR/PSD-95/SynGAP does not change at all (Figure Appendix B.3).

## 4. Discussion

### 4.1 Justification of our approach

Our model of only seven PSD proteins and without any specific spatial organization is definitely a highly simplified one that is far from the actual biological complexity of the postsynapse. In addition, for simplicity, we consider a situation that corresponds to a homozygous scenario, i.e. where either only wild-type or mutant Shank1 is present but not both. Last but not least, we have modeled only one well-defined effect of the mutation, ignoring possible pleiotropic effects like the alteration of the expression level of multiple proteins as observed for several Shank mutations [28]. Thus, it is not expected that the obtained protein complex distributions can be directly compared to the *in vivo* situations. Modeling all these aspects with acceptable accuracy would require much more data than currently available. However, we argue that our model system, focusing on a well-defined set of major PSD proteins and interactions is complex enough to capture general aspects of the behavior of elaborate protein networks with a multitude of binding interactions while remaining manageable in terms of data analysis as the number of possible protein complexes is not extremely high. On the other hand, mechanistic linking of genotypes with phenotypes, with different genotypes leading to similar phenotypes, is only possible *via* a combination of experimental data and modeling approaches.

### 4.2 Weakening a specific interaction can cause limited but significant changes

Mechanistic linking mutations to the phenotypes they cause is many times a non-trivial task, especially when the mutation perturbs a highly complex protein network. This phenomenon is well known in the case of neurodevelopmental disorders, where similar observed phenotypes can be caused by a number of different mutations. For example, a recent recommendation for Phelan-McDermid syndrome puts emphasis on the underlying genetic cause as the phenotypes are largely non-specific and can generally occur in a number of neurodevelopmental diseases [11].

The specific effect of mutations can be enigmatic, especially when they affect proteins present in many different tissues. This is especially true for cell types in which even the major partners and interactions are expected to be the same. The diversity of neurons in terms of the different abundance of postsynaptic proteins offers a unique opportunity to explore the effect of specific mutations in a complex but still simplified multicomponent system having the same set of building blocks. Our simulation-based approach, focusing on the formation of protein complexes as defined by the abundance of their constituent proteins and their interactions, reveals that the effect of a mutation weakening a specific interaction heavily depends on the availability of all interaction partners in the system. The complex interdependence of the interactions leads to a scenario where the overall changes in protein complex distributions are generally subtle, and the formation of only a few complexes are significantly affected and this effect can be confined to a well-defined set of cells with specific protein abundance. While common wisdom could suggest that protein complexes containing the mutated proteins are most affected and in cells where these are abundant, our results indicate that cells with the lowest number of affected proteins can also be among the vulnerable ones, and the protein associates mostly affected are linked only indirectly to the actually weakened interaction. Our simulations suggest that the cerebellum may be an involved brain region, aligning with findings in the literature [26, 25, 27].

Our results also suggest that mutations that (also) affect protein expression levels may have a more serious effect than those that moderately perturb a given interaction. In addition, the effect of redundancy in the system (e.g. Shank1-Shank3) might not only cause individual isoforms directly take over each other’s roles, but rather they reduce intermediary effects, e.g. here, the change in the frequency of the most important complex would always be moderated by the presence of a protein with a binding pattern similar to that of the mutant.

## 5. Acknowledgments

This research was supported by a grant from the National Innovation, Research and Development Office th the grant OTKA 137947.

## Appendix A. Supplementary Tables

*Appendix A.1. Table A: Input Proteins with Uniprot ID and Domains Appendix A.2. Table B: Domain-domain interactions*

*Appendix A.3. Table C: Brain regions*

*Appendix A.4. Table D: Function of brain regions*

*Appendix A.5. Table E: Input protein abundance data as used in the simulations*

*Appendix A.6. Table F: Ordered p-values of the complex AMPAR/PSD95/SynGAP*

*Appendix A.7. Table G: Ordered averaged p-values of the regions Appendix A.8. Table H: Wild-Type PCA coordinates*

*Appendix A.9. Table I: mutant PCA coordinates*

*Appendix A.10. Table J: wild-type and mutant PCA coordinates Appendix A.11. Table K: input abundance PCA coordinates*

*Appendix A.12. Table L: equilibrium dissociation constants found in literature*

## Appendix B. Supplementary Figures

*Appendix B.1. Figure A:* ***The abundances of complexes in the regions:***

A) H376.XI.50_CBC wild-type B) H376.XI.50_CBC mutant

C) H376.XI.52_CBC wild-type D) H376.XI.52_CBC mutant.

*Appendix B.2. Figure B:* ***The abundances of complexes in the regions:***

A) H376.IIIB.53_M1C-S1C wild-type B) H376.IIIB.53_M1C-S1C mutant

C) H376.IV.54_STR wild-type D) H376.IV.54_STR mutant. The complexes are: 0:NMDAR/PSD-95;1:NMDAR/PSD-95/SYNGAP;

4:PSD-95/AMPAR;5:PSD-95/AMPAR/SYNGAP;

6:PSD-95/AMPAR/SYNGAP/GKAP;8:PSD-95/SYNGAP;

10:PSD-95/GKAP;218239:GKAP/Shank1;

221654:GKAP/Shank1/Homer1;222783:Homer1-tetramer

*Appendix B.3. Figure C:* ***The abundances of complexes in the regions:***

A) H376.XI.50_HIP wild-type B) H376.XI.50_HIP mutant

C) H376.VIII.53_MD wild-type D) H376.VIII.53_MD mutant.

The complexes are: 0:NMDAR/PSD-95;1:NMDAR/PSD-95/SYNGAP; 4:PSD-95/AMPAR;5:PSD-95/AMPAR/SYNGAP;

6:PSD-95/AMPAR/SYNGAP/GKAP;

8:PSD-95/SYNGAP;10:PSD-95/GKAP;218239:GKAP/Shank1;

218240;218242 and 218243: variations of GKAP/Shank1/SHank1; 221654:GKAP/Shank1/Homer1;222783:Homer1-tetramer

*Appendix B.4. Figure D:* ***p-values plotted on brain***

*Appendix B.5. Description A: NeEMO*

Specifically, the NeEMO method uses sequence and structure information which is transformed into a residue interaction network. The residue interaction network is further processed for example by the Dijkstra algorithm and generates input data for a trained neural network that predicts the ΔΔ*G* for a given mutation.

The ΔΔ*G* refers to the entire protein, not domain-specific. As a result, the disadvantage is that the changes in all bonds of a given protein are treated equally and only estimated from the degree of change in stability.The change rate of the unfolding rate 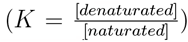 can be derived from ΔΔ*G*. The change rate tells how the unfolding rate changes by the mutation according to the predicted ΔΔ*G* values assuming that the Δ*G* is in equilibrium.

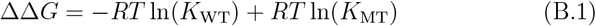

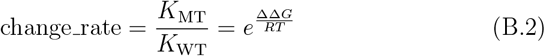

The probability of complex formation is inversely proportional to the change of unfolding rate in our model thus the change rate shows how much the unbinding rate changes.

