## Supplementary Figures for "Simulated complexes formed from a set of postsynaptic proteins suggest a localised effect of a hypomorphic Shank mutation"

 (ACSN)Attila Csikász-Nagy (ZG)Zoltán Gáspári inst1a inst2b

### Supplementary Figures

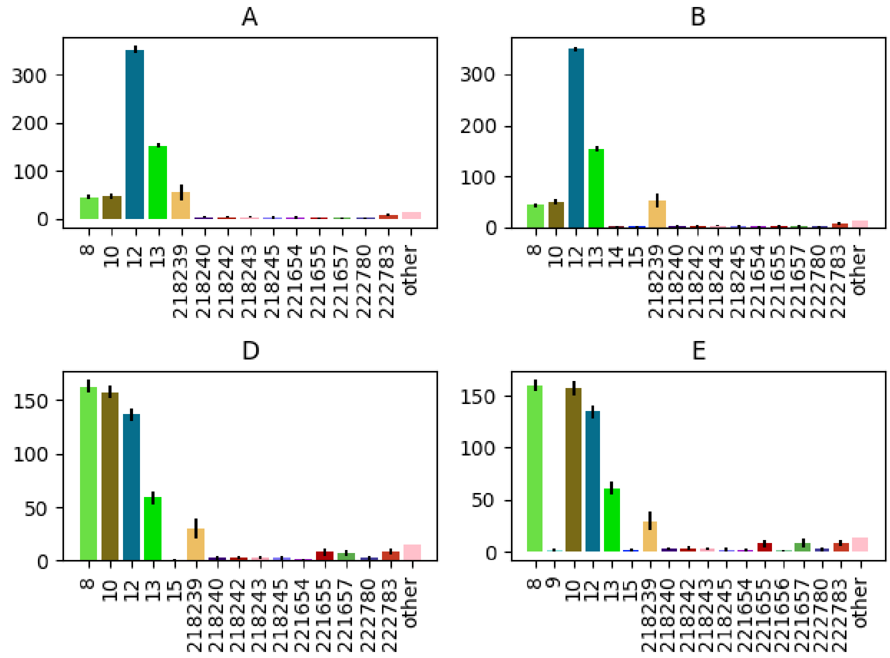


Figure S1: **The abundances of complexes in the regions:** A) H376.XI.50_CBC wild-type B) H376.XI.50_CBC mutant C) H376.XI.52_CBC wild-type D) H376.XI.52_CBC mutant.

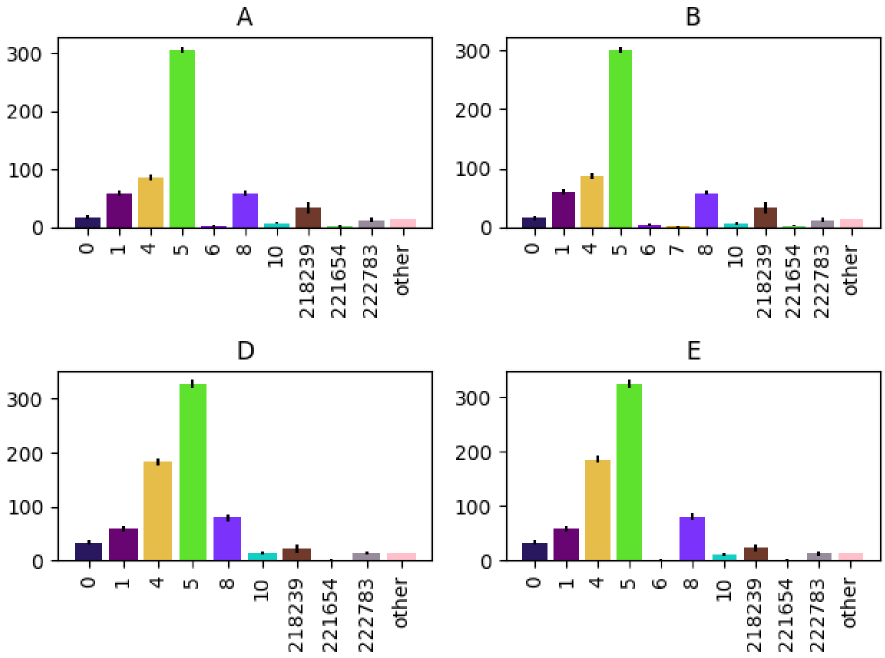


Figure S2: **The abundances of complexes in the regions:** A) H376.IIIB.53_M1C-S1C wild-type B) H376.IIIB.53_M1C-S1C mutant C) H376.IV.54_STR wild-type D) H376.IV.54_STR mutant. The complexes are: 0:NMDAR/PSD-95;1:NMDAR/PSD-95/SYNGAP;4:PSD-95/AMPAR;5:PSD-95/AMPAR/SYNGAP;6:PSD-95/AMPAR/SYNGAP/GKAP;8:PSD-95/SYNGAP;10:PSD-95/GKAP;218239:GKAP/SHank1;221654:GKAP/SHank1/Homer1;222783:Homer1-tetramer

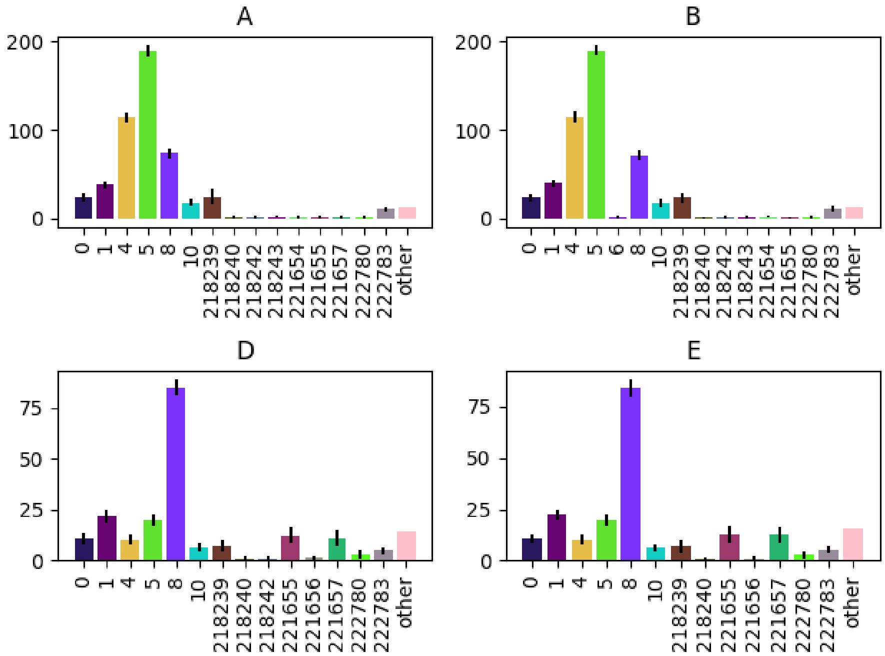


Figure S3: **The abundances of complexes in the regions:** A) H376.XI.50_HIP wild-type B) H376.XI.50_HIP mutant C) H376.VIII.53_MD wild-type D) H376.VIII.53_MD mutant. The complexes are: 0:NMDAR/PSD-95;1:NMDAR/PSD-95/SYNGAP; 4:PSD-95/AMPAR;5:PSD-95/AMPAR/SYNGAP;6:PSD-95/AMPAR/SYNGAP/GKAP;8:PSD-95/SYNGAP;10:PSD-95/GKAP;218239:GKAP/SHank1;218240;218242 and 218243: variations of GKAP/SHank1/SHank1;221654:GKAP/SHank1/Homer1;222783:Homer1-tetramer

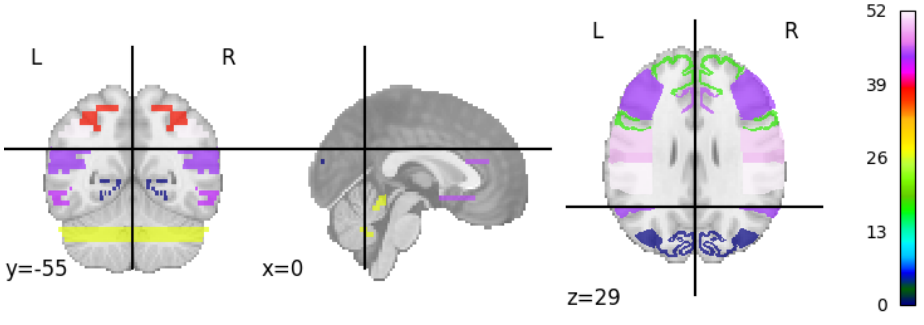


Figure S4: **p-values plotted on brain**
